# New insights in host-associated microbial diversity with broad and accurate taxonomic resolution

**DOI:** 10.1101/050005

**Authors:** Matthew T. Agler, Alfredo Mari, Nina Dombrowski, Stéphane Haquard, Eric M. Kemen

## Abstract

- Deep microbiome profiling has sparked a revolution in biology, recontextualizing mechanisms such as macroorganismal adaptation and evolution. Amplicon sequencing has been critical for characterization of highly diverse microbiomes, but several challenges still hinder their investigation: (1) Poor coverage of the full diversity, (2) Read depth losses and (3) Erroneous diversity inflation/distortion.
- We developed a modular approach to quickly profile at least 8 interchangeable loci in a single sequencing run, including a simple and cost-effective way to block amplification of non-targets (like host DNA). We further correct observed distortion in amplified diversity by phylogenetically grouping erroneous OTUs, creating a phylogeny-based unit we call OPUs.
- Our approach achieves full, accurate characterization of a mixed-kingdom mock community of bacteria, fungi and oomycetes at high depth even in non-target contaminated systems. The OPU concept enables much more accurate estimations of alpha and beta diversity trends than OTUs and overcomes disagreements between studies caused by methodology. Leveraging the approach in the *Arabidopsis thaliana* phyllosphere, we generated to our knowledge the most complete microbiome survey to date.
- Microbiomes are extremely diverse, extending well beyond bacteria and fungi. Our method makes new questions in a variety of fields tractable with accurate, systems-based overviews of microbial community structures.

## Introduction

A revolution in biology is currently underway as our understanding of various systems is brought into the context of newly characterized structures and roles of symbiotic microbial consortia. This transformation is the result of growing research on microbiota associated with various abiotic or biotic systems (270 vs. 3494 publications had the words “Microbial community” or “Microbiome” in the title in 2005 vs. 2015 according to a PubMed search on Mar 22, 2016). Strong interest in this field is not surprising considering that research is turning up important roles of the community context of microorganisms in systems as diverse as biotechnological transformations (Werner *et al.*, 2011) and plant and animal health and fitness (Hehemann *et al.*, 2010, Mills *et al.*, 2013, Panke-Buisse *et al.*, 2015, Rolli *et al.*, 2015).

A typical approach employed by microbiome researchers is first to characterize microbial community structures in a system of interest. To do so, many studies rely on amplicon sequencing of phylogenetically informative genomic loci to generate microbiota profiles. These profiles are then linked to specific experimental parameters, host phenotypes or performance measurements. Community profiling based on ribosomal gene phylogeny dates to Pace and colleagues (Stahl *et al.*, 1985) who in 1985 reported on isolating and sequencing the 5S rRNA gene in environmental samples to identify abundant but uncultured bacteria. The technology has come a long way: As with many recent important developments in biology, rapid and inexpensive DNA sequencing technology has been an enabling force in microbiome research. Its democratization, however, is due to development of highly parallel library indexing which made high-throughput amplicon sequencing extremely inexpensive on a per sample basis (Hamady *et al.*, 2008).

Today, with for example the MiSeq platform, amplicon libraries are routinely and rapidly generated from hundreds of samples and sequenced together in a single run. This process generates millions of sequences up to 600 bp in length (Caporaso *et al.*, 2012), enabling extremely deep profiling of targeted microbial groups. In the first experiment of the current study, we used the Illumina MiSeq to characterize *A. thaliana* root compartments more deeply than we could previously with 454 pyrosequencing (Schlaeppi *et al.*, 2014) in hope of gaining new insights. We show that better diversity recovery with the Illumina protocol, not read depth, enabled better differentiation of soil and rhizosphere compartments. In addition to the need to maximize microbial diversity coverage, we identified two other problems limiting characterization of diverse microbial communities: (1) Losses to read depth because of non-target amplification and (2) Artificial inflation/distortion of diversity due to erroneous OTUs.

Limited profiling of diversity extends well beyond bacteria, since microbiomes are often composed of species from all kingdoms of life. These cohabiting members interact with the environment and influence one another via direct associations (Fisher & Mehta, 2014) or indirectly via a host (Hajishengallis, 2015). To resolve these interactions and model microbial community dynamics, robust systems approaches are needed (Lima-Mendez *et al.*, 2015). For example, analysis of modularity in microbial correlation networks (i.e., co-occurring groups of microbes) has revealed rice root-associated prokaryotes involved in methane cycling (Edwards *et al.*, 2015) as well as modules of fungi and bacteria that together correlate with certain soil parameters (de Menezes *et al.*, 2015). To improve the usefulness of such approaches, some studies are profiling a larger diversity like bacteria and fungi simultaneously (Marupakula *et al.*, 2016). Such approaches can reveal, for example, keystone species that underlie microbial community structures because they interact heavily, linking external abiotic and biotic sources of variation to the community (Berry & Widder, 2014). Recent studies in phyllosphere microbial communities (Agler *et al.*, 2016) and in ocean samples (Chow *et al.*, 2014) have emphasized that keystone microbes participate heavily in inter-kingdom interactions. Thus, broad coverage of diversity is critical to pinpoint these important microbes in community surveys.

Parallel amplification and sequencing of multiple loci is one way to cover more diversity and for this approach many well-characterized targets are available. Among other target loci, structures of communities can be probed *via* the 16S rRNA gene (Baker *et al.*, 2003), the internal transcribed spacer (ITS) region 1 or 2 (Blaalid *et al.*, 2013) or the 18S rRNA gene (Hugerth *et al.*, 2014) for prokaryotic, fungal, and other eukaryotic microbes, respectively. For most of these targets, many possible primer sets are available, each bringing their own biases and specificities. Therefore, including multiple loci from a single gene target can be advantageous and provide complementary information (Wang *et al.*, 2016). Whatever the target choice, modular methods are needed to quickly adapt methodology to specific questions because represented diversity varies considerably between different microbial communities. Previously, we developed a method to prepare, sequence and analyze two loci from each of bacteria (16S), fungi (ITS) and oomycetes (ITS) in parallel and used it to evaluate microbial structure and interactions in the *Arabidopsis thaliana* phyllosphere (Agler *et al*, 2016). Here, we use a mixed mock community of microorganisms to optimize it for high throughput, resolution and accuracy.

We also address the two other major barriers to full characterization of microbial communities. The first is amplification of host or non-target organismal DNA such as mitochondrial 16S, chloroplast 16S or non-target genomic ITS sequences that can lead to major loss of useful reads (Bulgarelli *et al.*, 2012, Ihrmark *et al.*, 2012). We developed “blocking oligos” that inexpensively and nearly completely eliminated non-target amplification in mock communities with simulated “host” contamination. We also show that their use does not bias results and that they can be adapted to block amplification of undesirable microbial targets. Second, we addressed false trends in recovered microbial community diversity arising because of sequencing errors (Kunin *et al.*, 2010). Here, we introduce the “operational phylogenetic unit” (OPU) – phylogenetic groupings of erroneous OTUs. We show that this method resolves differences between our 454 and Illumina methods caused by errors. We also demonstrate that OPUs can be used to generate a phylogenetic beta diversity distance metric even for fungal ITS reads and that they are a much more accurate direct measure of species richness than OTUs.

Finally, we leverage all of these benefits to profile microbes associated with leaves of wild *A. thaliana* plants. To demonstrate full modularity and that any desirable locus could be included, we expand to target microbial eukaryotes with two loci of the 18S rRNA gene (8 loci total). The result is deep profiling of all 8 loci and to our knowledge the most complete picture to date of diversity in a microbial community. We provide all tools and information needed for researchers to analyze up to 50 samples with the 8 loci described here or to expand the system for their needs. Together, these simple solutions will enable researchers to rapidly, accurately and nearly completely characterize many microbial communities in a single sequencing run. We expect these methods to broaden the applicability and impact of amplicon sequencing experiments.

## Materials and Methods

### Comparison of methods for amplicon sequencing

We first tested an amplification and sequencing protocol for the Illumina MiSeq that includes amplicon generation in two PCR steps (**Fig. S1**). We prepared bacterial 16S rRNA gene libraries from 3 bulk soil, 3 plant rhizosphere and 2 plant root samples from the ‘Eifel’ natural site experiment (Experiment 1, **Table S1**) and sequenced them as described in **SI methods**. We combined Illumina data with data from the same samples previously generated using 454 technology (Schlaeppi *et al*, 2014) and generated OTUs as described in **SI methods**. We then summarized OTUs by taxonomy and generated plots at the phylum level (all taxa) or the family level (the 20 most abundant taxa). We compared the cumulative OTU discovery vs. depth between the two technologies by rarifying tables at read depth intervals of 10 between 0 and 100 and 100 between 100 and 3000 and counting the number of unique and shared OTUs generated by each technology. Finally, we compared the ability of the technologies to discriminate between compartments of *Arabidopsis thaliana* roots by plotting boxplots of Bray-Curtis or weighted UniFrac distances between sample classes using phyloseq (McMurdie & Holmes, 2013) and ggplots2.

### Optimizing modular, multi-locus library preparation and sequencing

We expanded the method used in the first experiment to target multiple loci in a single sequencing run (Experiment 2, **Table S1** and **Table S2**). Accuracy was tested by amplifying mixed kingdom mock communities (**Table S3**) in 6 separate PCR reactions targeting two loci from phylogenetically informative regions of each of bacteria (16S rRNA V3-V4 and V5-V7), fungi (ITS1 and 2) and oomycetes (ITS1 and 2). We tested effects of library preparation methodology by performing PCR in one step (35 cycles) or two steps (10 then 25 cycles or 25 then 10 cycles) (**Fig. S1b** and **Table S1**). For two-step preparations, the primers used in the first step consisted of unmodified universal amplification primers (**Fig. 1a**). For single-step preparations and for the second step in two-step preparations, primers were a concatenation of the Illumina adapter P5 (forward) or P7 (reverse), an index sequence (reverse only), a linker region, and the universal primer for the region being amplified (**Fig. 1b**, **Fig. S1a** and **File S1**). Details of all PCR steps can be found in the **SI methods**. Libraries were purified, quantified, and combined in equimolar concentrations. Sequencing was on a single Illumina MiSeq lane (Illumina, Inc.) by adapting the approach of Caporaso et al. (Caporaso *et al*, 2012) for multiple loci (**Fig. S1c** and **File S1**). This recovers ~8% more high quality bases than protocols relying on Illumina sequencing primers (calculated in **SI Note**).

**Figure 1.**
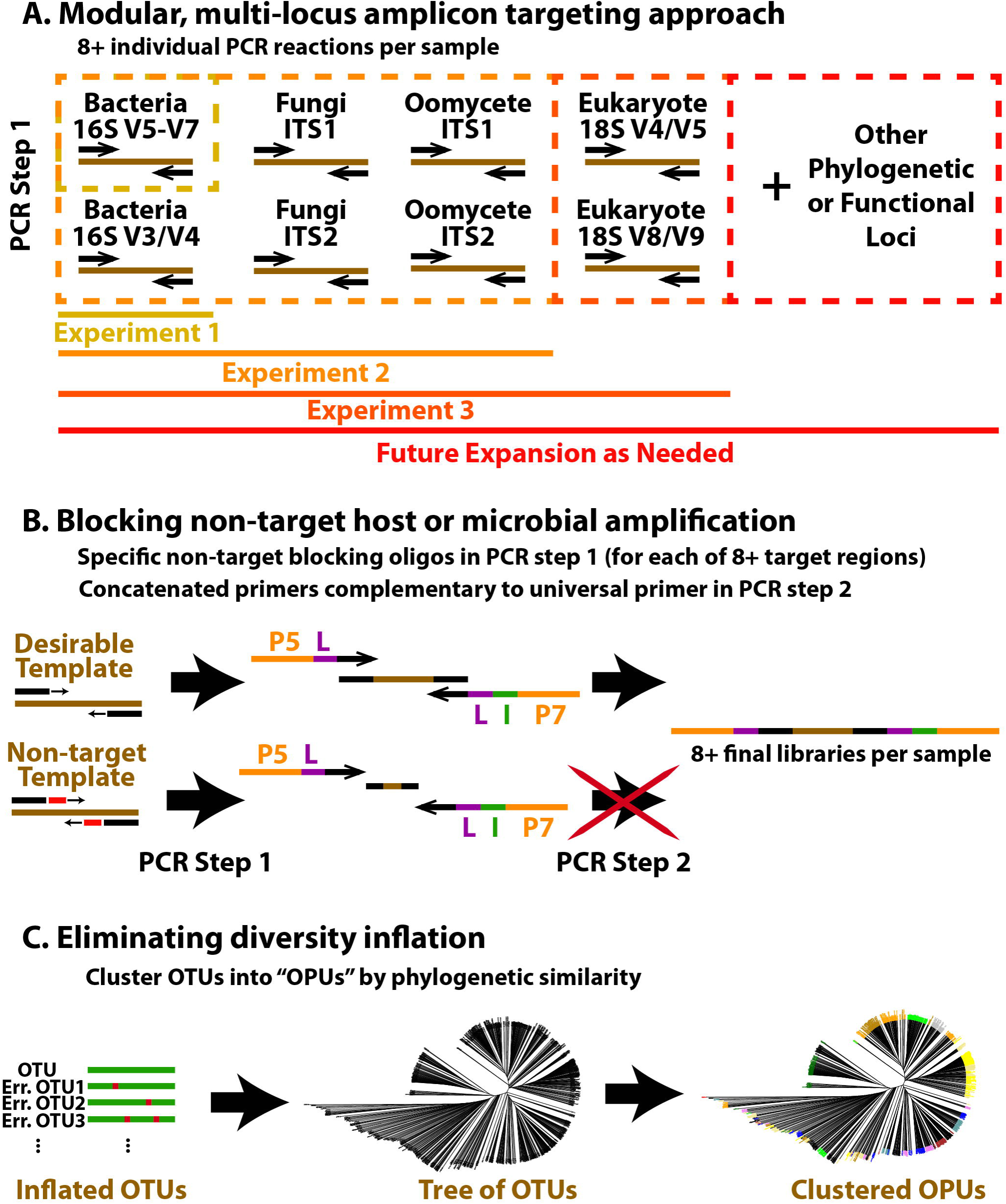
Strategy to increase taxonomic coverage and accuracy of amplicon sequencing. A. In the first step, 8 individual PCR reactions are performed per sample each targeting a specific gene region. B. Blocking oligos are employed in the first PCR step which are specific for non-target templates so that these cannot be elongated to the final libraries with concatenated primers in the second step. P5 and P7 are standard Illumina adapter sequences. L indicates the linker sequence. C. Inflation of the number of observed OTUs caused largely by sequencing error is addressed by dividing them up by taxonomic lineages, building a phylogenetic tree and then clustering closely related members into operational phylogenetic units (OPUs).

Details on generating OTU tables and taxonomy from raw multi-locus data can be found in the **SI Methods**. We summarized bacterial, fungal and oomycete OTU tables by taxonomic rank, converted abundances to relative values and plotted the family-level taxonomic distribution directly from this data with the package ggplots2 in R. To calculate distances of samples from expected, we added the expected distributions (**Table S3**) to the OTU tables and summarized taxa at the family level. After removing “host”-derived reads, we calculated Bray-Curtis distances between samples using Vegan (Oksanen *et al.*, 2013). We plotted distances from expected distributions in boxplots using ggplots2. Each box represents three “replicate” libraries generated with the same mixed kingdom mock community template but with differing amounts of “host” DNA added.

### Avoiding non-target template amplification with “blocking oligos”

To make the method applicable to host-associated studies, we addressed non-target amplification in library preparation. In short, primers specific to the known, undesirable template (hereafter “blocking oligos”) are designed to bind nested inside the universal primer binding sites (Experiment 3, **Table S1**). Thereby, most amplicons made in the first PCR step from non-target template are short and lack the universal primer sequences. These cannot be elongated in the second PCR step and subsequently are not sequenced (**Fig. 1b**, **Fig. S1b** and **S1c**). Blocking oligos were designed for the *A. thaliana* chloroplast (16S rRNA V3-V4 region) or mitochondria (16S rRNA V5-V7 region) and the *A. thaliana* ITS1 and ITS2 regions by adapting the approach of Lundberg et al. (Lundberg *et al.*, 2013) originally for PNA clamps. See **SI Methods** and **Fig. S2** for details of design and use in library preparation and **File S1** their sequences. To analyze the percent reduction in host plant-associated reads when blocking oligos were employed, we considered the relative abundance of reads associated with the class “Chloroplast” or the order “*Rickettsiales*” in the 16S OTU tables and reads in the kingdom “*Viridiplantae*” in the ITS OTU tables in samples with *A. thaliana* DNA and with and without blocking oligos.

### Clustering OTUs by phylogeny into OPUs

For the 454/Illumina comparison and multi-kingdom mock community data, OTUs were clustered into phylogenetically closely related groups that we called operational phylogenetic units (OPUs, **Fig. 1c**, Experiment 4 in **Table S1**). In short, OTUs were divided at the rank of family, combined with sequences from the taxonomy reference databases and a phylogenetic tree was built for each by alignment with MUSCLE (Edgar, 2004). UPGMA trees for each family were created with the R function hclust. The tree was dynamically split into clusters using the hybrid method in cutreeDynamic in the dynamicTreeCut package (Langfelder *et al.*, 2008) in R. This method was designed to identify clusters in trees similar to hierarchical clustering but without predetermined clustering depths. It dynamically identifies groups of tips in a dendrogram that form clusters using both the tree and the distance matrix that the tree is based on. A set of user-defined parameters define the cluster detection sensitivity and we found that setting the minimum cluster size to 15 and the deepSplit parameter to 3 was effective for OTU clustering. We then generated a map of the OTUs in each OPU and generated an OPU abundance table.

For the 454/Illumina data, overlap of OPU generation between technologies and Bray-Curtis distance plots were generated exactly as described above for OTUs. For mock communities, species richness estimates were based on data from the evenly distributed mock community template with *A. thaliana* “host” contamination, amplified in 2 steps (10 cycle / 25 cycle) with blocking oligos. We used QIIME 1.8.0 (Caporaso *et al.*, 2010) to calculate the number of observed species in 10 rarefactions at 30 evenly spaced depths based on the OPU table, the OTU table, and the tables of OTUs grouped taxonomically at levels species, genus, family and order. The maximum depth was based on the OPU read depth since a few reads were discarded during OPU generation (Bacteria V3/V4: 2530, V5/V6/V7: 34780, Fungi ITS1: 9930, ITS2: 48400, Oomycete ITS1: 26820, ITS2: 5000). We plotted the average number of observed species against sequencing depth for the bacterial 16S V3-V4 dataset and the ratios of observed:expected OTUs and OPUs for all datasets.

### Characterizing A. thaliana phyllosphere microbiota

We used the multi-locus approach to characterize the phyllosphere microbiome of *A. thaliana* leaves infected with the oomycete pathogen *Albugo laibachii* with near-complete taxonomic coverage (Experiment 5 in **Table S1** and **Table S2**). Whole leaves (defined as a single whole rosette) or endophytic fractions of leaves (defined as in (Agler *et al*, 2016)) were collected in the wild (a total of 18 samples - 9 whole leaf, 9 endophyte) and were immediately frozen on dry ice. DNA extraction was performed as described previously (Agler *et al*, 2016). Library preparation, sequencing and analysis was performed as described above. To more completely cover eukaryotic microbial diversity, we expanded the 6 loci method to 8 with two additional 18S rRNA gene loci (V4-V5 and V8-V9, see **File S1**).

To reduce *A. thaliana* or *A. laibachii* amplification in the 18S region we designed additional blocking oligos for both of these organisms (**File S1**). We tested them by preparing 18S amplicons from two mock communities consisting of *A. thaliana* (97% or 87%), *A. laibachii* (0 or 10%), *Sphingomonas* sp. (1.5%), *Bacillus* sp. (1.5%) and 0.001% to 1% of target *Saccharomyces* cerevisiae. (**Table S4**).

To provide a complete and concise picture of the diversity of microbiota inhabiting *A. thaliana*, we combined the data from all samples. To visualize data, we assigned taxonomy to OTUs and generated two phylogenetic trees where branches represent unique genera. Trees were generated from the taxonomic lineages (*not* OTU sequence similarity) with the ape package in R and output as newick files (Paradis *et al.*, 2004). Therefore, OTUs from taxa not represented in the databases are simply grouped as “Unclassified”. These were uploaded to iTOl v3.1 (Letunic & Bork, 2016) to color branches by taxonomy or by targeted regions. The first tree, for Eukaryotes, includes data from the 18S and ITS targeted regions. The second tree includes data from the 16S targeted regions.

### Data Availability and Figure Regeneration

Raw sequencing data is being made publicly available *via* Qiita (https://qiita.ucsd.edu/) study number 10408 and is currently available for direct download at: http://bioinfo.mpipz.mpg.de/download/MethodPaper_Share/. All modified databases, OTU tables and metadata files, as well as scripts and instructions to generate OPUs and recreate the main figures are available at Figshare (https://figshare.com/s/07b3493d1f6442d34dfd).

## Results

### Pattern recovery depends on diversity coverage but is obscured by erroneous OTUs

We first re-analyzed the 454-generated bacterial 16S data from (Schlaeppi *et al*, 2014), confirming that rhizosphere and soil compartments from *A. thaliana* roots were weakly distinct (**Fig. 2a**). We hypothesized that because of their relatively high alpha diversity, higher read depth was needed to differentiate microbiota between the compartments. Thus, we reanalyzed the same set of samples at higher depth (86,406-211,907 reads/sample vs. 12,699-20,844 reads/sample previously) with our protocol for the Illumina platform (**Fig. 1a and 1b and Table S1**). Bray-Curtis distances, which consider all OTUs equally, suggested that the Illumina method indeed better distinguished rhizosphere and soil compartments (**Fig. 2a**). Surprisingly, this was depth-independent and was also true for differences between other compartments. This was apparently driven by widely divergent OTU profiles with only 35% of all OTUs observed in both datasets (**Fig. 2b**, 700 and 1424 OTUs were unique to 454 and Illumina, respectively). Huge numbers of unique OTUs suggested that either: (1) Differences in the methods of library generation and sequencing resulted in little overlap or (2) OTUs were inflated by error. To check this, we calculated between-sample weighted UniFrac distances, which gives less importance to differences caused by closely related, likely erroneous OTUs (**Fig. 2a**). With this metric only soil and rhizosphere compartments were better differentiated in the Illumina dataset, suggesting a mixture of real differences and erroneous OTUs leading to false diversity trends. True differences could be due to higher sensitivity with the Illumina method to the phyla *Verrucomicrobia*, *TM7* and *Chloroflexi*, which were more abundant in that dataset (**Fig. S3**). However, there were apparently too many errors to locate soil/rhizosphere differential OTUs with certainty. In any case, the role of improved taxonomic resolution in detecting fine differences between datasets motivated us to expand to target loci beyond prokaryotes.

**Figure 2.**
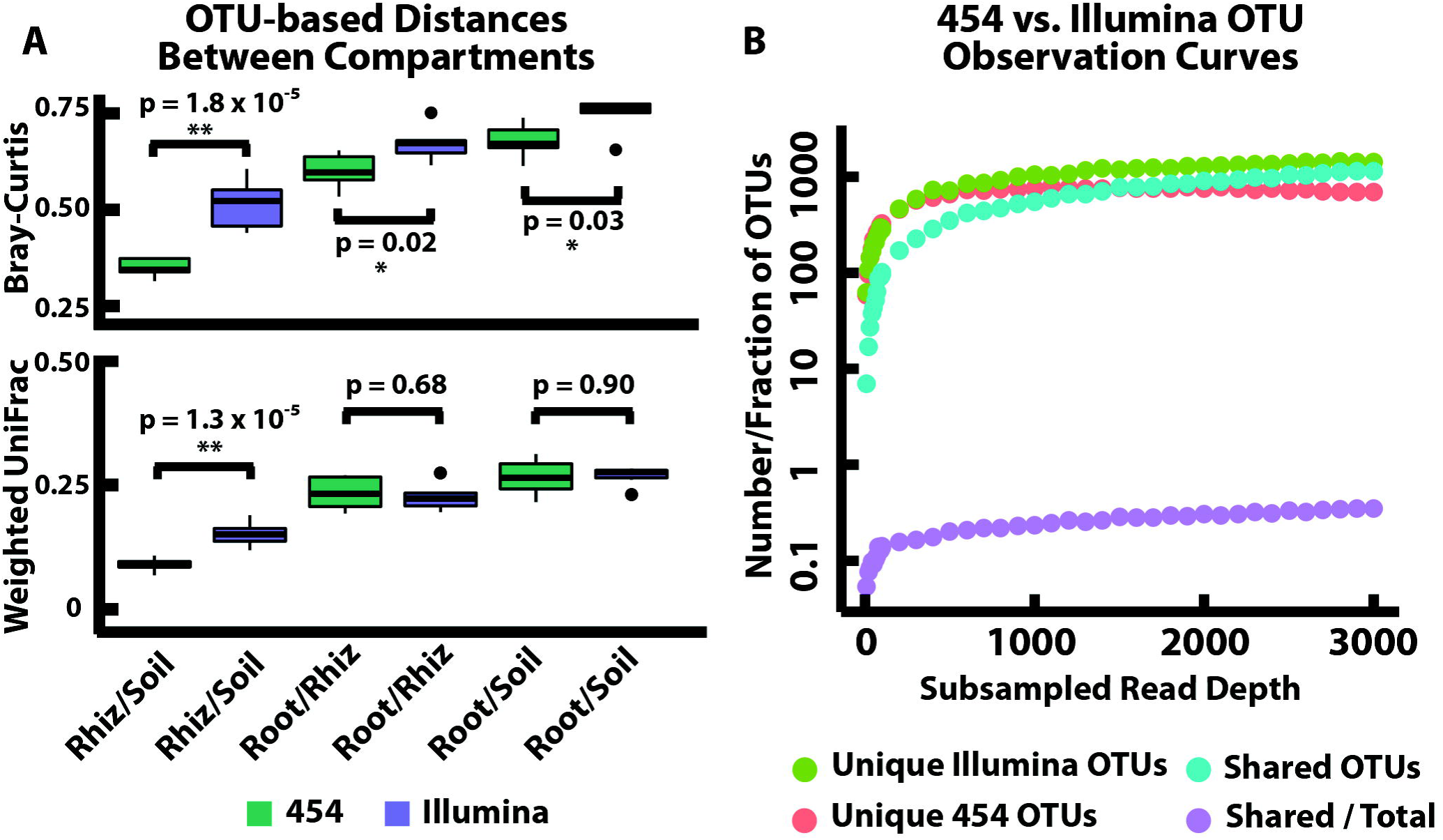
A comparison of 454- vs. Illumina-based amplicon sequencing protocols shows little overlap of OTUs, suggesting high erroneous OTU generation. A. Bray-Curtis distances based on OTU relative abundances suggest that data recovered with the Illumina protocol significantly better distinguishes all pairs of compartments. Weighted UniFrac, however, suggests that the Illumina protocol only better distinguishes rhizosphere and soil compartments, implying that differences between others were due to closely related and probably erroneous OTUs. B. Between 454 and Illumina datasets, rarefaction curves of the number of OTUs discovered with increasing read depth suggest that erroneous OTUs are very common since most OTUs are unique to datasets produced by one or the other technology, with only about one-third of all OTUs found in both datasets. The rarefaction curves are separated into OTUs that are unique to Illumina, unique to 454 or shared by both technologies.

### A fully modular, multi-locus approach to improve insight into microbial diversity

We previously (Agler *et al*, 2016) adapted our 2-step Illumina amplicon library generation protocol to simultaneously target 6 genomic loci, two from each of bacteria, fungi and oomycetes. Here, we optimized the protocol by extensively testing variations of it on a mock community consisting of microbes from the three kingdoms (**Table S3**). We found that for all three kingdoms, taxa distributions (shown at the order level in **Fig. 3a** **and Fig. S4a**) were similar to expected. Mocks with staggered distributions of microorganisms were generally closest to expected (**Fig. 3b** **and Fig. S4b**) because the effect of underestimated taxa was sometimes stronger in even communities (e.g., the order *Mucorales* was not efficiently recovered by fungal ITS1 primers **Fig. 3a** **and Tables S5-S7**). Recovered community structures were reproducible, since the distance of technical replicates from the expected distribution was consistent (**Fig. 3b** **and Fig. S4b**). 2-step amplification recovered microbial community structures that were closer to the expected than 1-step amplification although the trend was not significant in all datasets. Further, leaving the bulk of PCR cycling for the second step (10 cycles followed by 25 cycles) tended to give the most accurate results. These close-to-expected taxonomy distributions were based on OTUs grouped by taxonomy. At the OTU level we again observed inflated diversity due to erroneous OTUs. OTUs overestimated species richness by on average 257.5%, 2575% and 387.5% (only considering OTUs in the expected taxa) for bacteria, fungi and oomycetes, respectively (**Tables S5-S7**).

**Figure 3.**
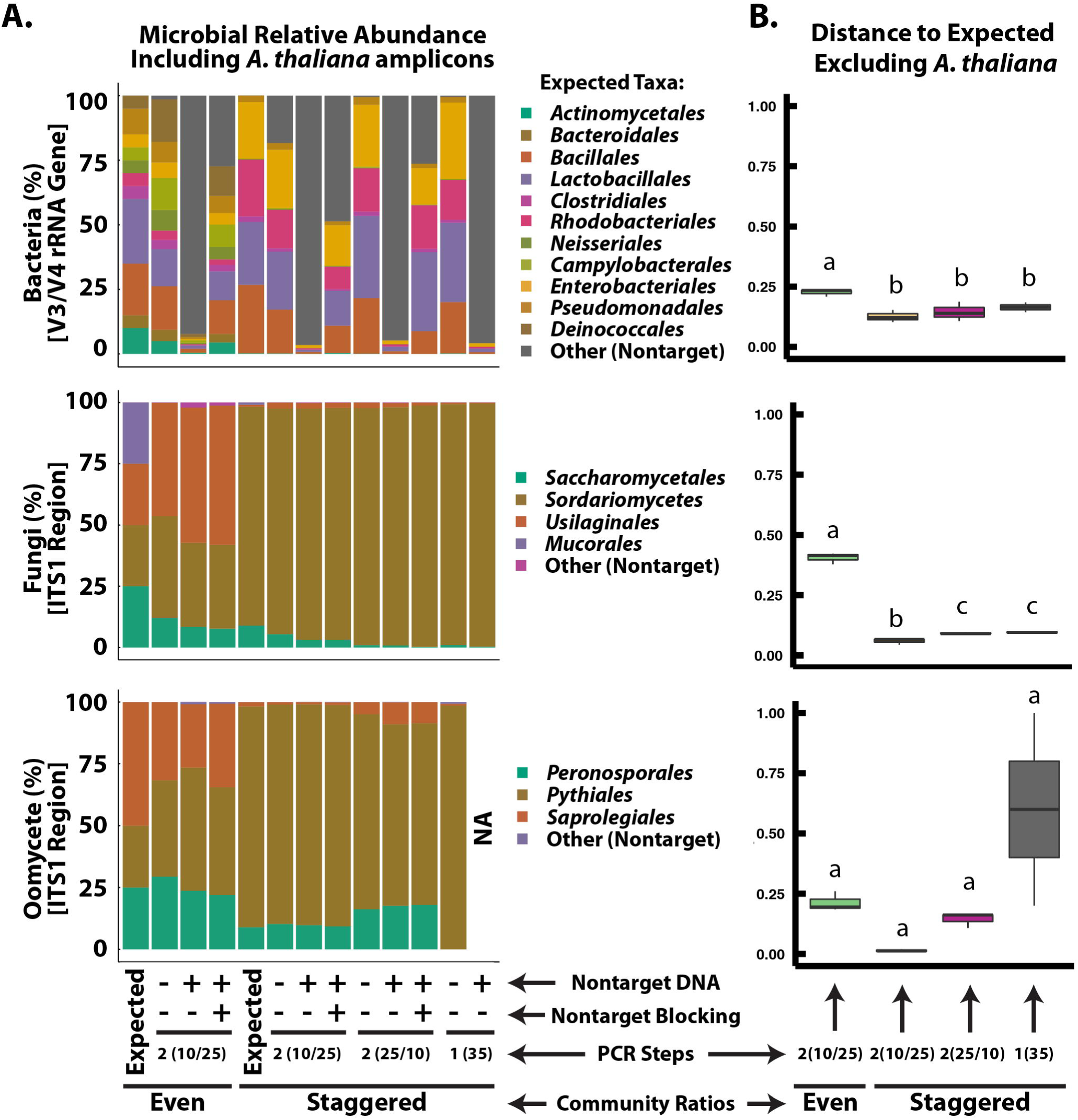
Reproducible and accurate characterization of mock communities of bacteria, fungi, and oomycetes by amplicon sequencing. A. Observed taxa at the order level in sequenced mock communities closely matched expected communities. The taxa “Other” is primarily non-target amplification from *A. thaliana* “host” DNA that was added to test blocking oligomers which prevent “host” DNA amplification. “NA” indicates a sample where sequencing depth was too low after subsampling to be included. B. Distance (Bray-Curtis distance based on relative abundance of order-level taxa) of sequenced communities from the expected distribution where 0 is identical and 1 is unrelated. Even or staggered revers to the distribution of the organisms in the mock communities (see expected distributions in A). PCR Steps refers to a 1-step (35 cycles with concatenated primers) or 2-step (10 or 25 cycles with standard primers followed by 25 or 10 cycles with extension primers containing Illumina adapters) amplification protocol. Letters indicate p < 0.1 (FDR-corrected) based on pairwise t-tests between groups.

We tested applicability of our method to host-associated microbiomes by mixing 90% *A. thaliana* “host” DNA and 10% mock communities (**Fig. 3**). Non-target host-derived DNA amplification accounted for up to 94% of reads (chloroplast-derived in the 16S V3-V4 dataset) and much less but still significant amounts in other target regions (**Fig. 3a** **and Fig. S4**). Therefore, we developed and implemented “blocking oligos” to reduce amplification of non-target DNA template. This method largely recovered read depth by eliminating 60 - 90% of chloroplast contamination in bacterial 16S communities and nearly all of the small amount of contamination in fungal ITS communities (**Fig. 4a**). Importantly, employing blocking oligos did not change the recovered distribution of taxa (each of the 2-step amplification boxplots in **Fig. 3b** included a replicate with blocking oligos but all had the same distance to expected). Thus, whereas extensive host contamination would obscure all but the most abundant microbes, blocking oligos enable deeper amplicon sequencing to uncover rare microbiota.

**Figure 4.**
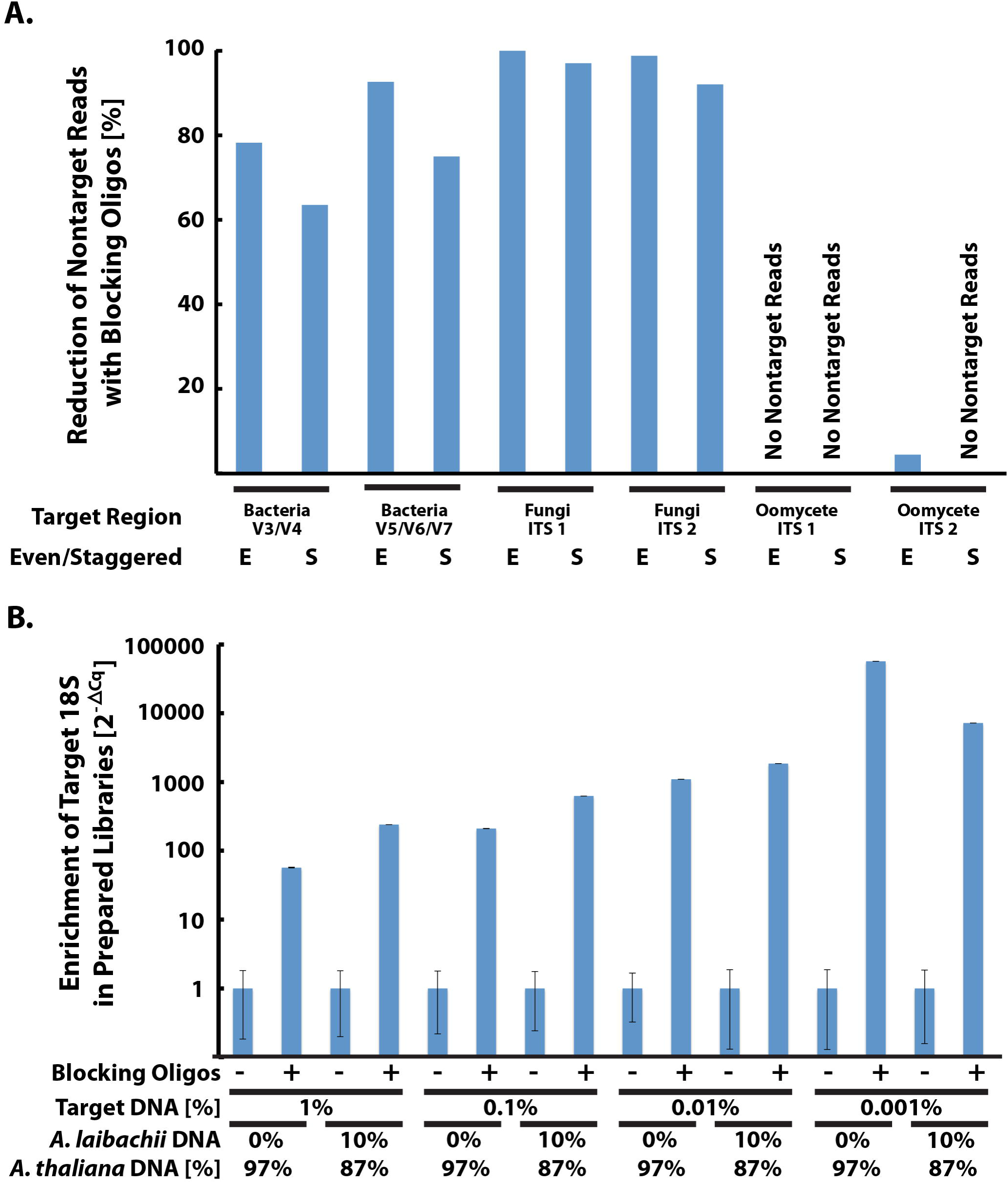
Employing blocking oligomers greatly reduces non-target and increases target yield in amplicon libraries. A. Near-complete reduction of amplification of *A. thaliana* “host” non-target plastid 16S or ITS by employing blocking oligos in preparation of mock community libraries. B. Relative increase of target (*Saccharomyces* sp.) 18S V4-V5 region amplicons (qPCR 2^-ΔCq^ values relative to measurement without blocking oligomers) in mock community libraries prepared with blocking oligomers to reduce *A. thaliana* and *A. laibachii* non-target amplification.

### Recognizing true diversity trends with phylogenetic OTU clustering

Prolific generation of erroneous OTUs strongly distorted true diversity patterns. Since erroneous OTUs derive from the same true sequence (**Fig. 1c**), they should cluster closely in phylogenetic trees. Using this principle we grouped OTUs generated in the 454/Illumina comparison into a unit that we call operational phylogenetic units (OPUs). This approach reduced the total from 3268 OTUs to 293 OPUs. OPUs properly grouped divergent erroneous OTUs generated with 454 and Illumina since overlap between them increased from 35% (OTUs, **Fig. 2b**) to 90% (OPUs, **Fig. 5a**). There were only 11 and 21 OPUs unique to 454- and Illumina, respectively. Bray-Curtis distances based on OPUs (**Fig. 5b**) closely resembled UniFrac distances based on OTUs (**Fig. 2a**) where Illumina only better distinguished soil and rhizosphere compartments. We determined that 16 OTUs (maximum 150.8 reads/sample) and 10 OPUs (maximum 213.7 reads/sample) significantly contributed to observed soil/rhizosphere differentiation (**Table S8**). Significant OTUs and OPUs were in agreement, since both were dominated by the phylum *Chloroflexi* (4 of 6 OTUs with > 20 reads/sample and the most abundant OPU, **Table S8**).

**Figure 5.**
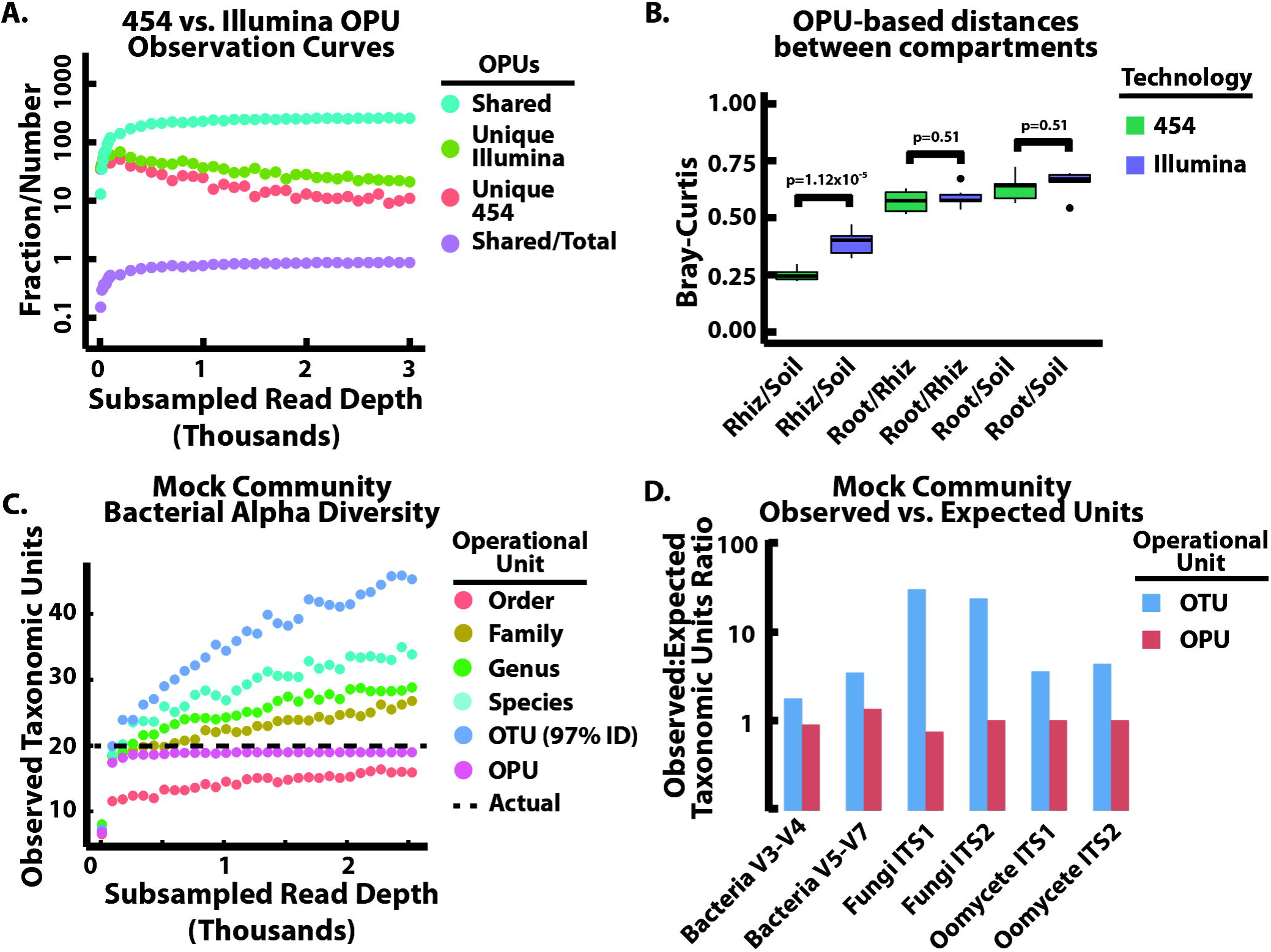
Clustering of operational taxonomic units (OTUs) into operational phylogenetic units (OPUs) by their phylogenetic relatedness corrects erroneous diversity discovery. A. Between 454 and Illumina datasets, the number of shared and unique OPUs and the fraction of shared OPUs demonstrates that OTU clustering greatly reduces erroneous dataset disagreements compared to Fig. 2B. B. Bray-Curtis distances based on OPUs displays similar trends as weighted UniFrac distances with the only significant differences between rhizosphere and soil compartments. C. Rarefaction curves of observed units at the OTU level, various taxonomic ranks, and for OPUs for bacterial 16S V3/V4 amplicon data show that unclustered OTUs and most taxonomic ranks greatly overestimate the expected diversity and the curves do not reach an asymptote, while OPUs quickly reach an asymptote close to the expected diversity. D. Numbers of observed OTUs or OPUs vs. expected units for all target regions demonstrates near-expected numbers of taxa in all target regions. Data in C. and D. is generated from the evenly distributed mock community with *A. thaliana* “host” DNA and using host-blocking oligomers and only considers OTUs and OPUs in the expected taxonomic families.

We used the mock community dataset to check if OPUs can provide accurate diversity estimates in all target loci. Indeed, all OPU rarefaction curves quickly reached an asymptote close to expected species richness (**Fig. 5c**). Considering only expected families, species richness was estimated at 110%, 87.5% and 87.5% of expected for bacteria, fungi and oomycetes, respectively (**Tables S5-S7**). For fungal and oomycete ITS2 data, richness was estimated perfectly in all families, and the same was true for most families in other datasets (**Tables S5-S7**). Considering all discovered OPUs and OTUs (including non-targets and contaminants), phylogenetic grouping reduced the average total number of observed units from 86 to 30.5 (OTUs to OPUs) for bacteria (20 expected), 121.5 to 7 for fungi (4 expected) and 26 to 7.5 for oomycetes (4 expected) (**Fig. 5d** **and Tables S5-S7**). Comparatively, OTUs and even OTUs grouped by taxonomy extensively overestimated diversity and their non-asymptotic rarefaction curve suggests continued inflation with deeper sequencing (**Fig. 5c**). Overall, OPUs contribute to drastically improved microbial diversity profiles from amplicon sequencing data.

### Towards a complete survey of complex host-associated microbiomes

Next, we leveraged the full modularity of our approach to provide a near-complete survey of prokaryotic and eukaryotic diversity in *A. thaliana* leaves collected in several locations using 8 loci (2 for each of bacteria, general eukaryotes, fungi and oomycetes). Since the leaves of *A. thaliana* are often infected by the obligate biotroph pathogen *A. laibachii*, templates from leaves are dominated by *Arabidopsis* and *Albugo* genomic DNA. To overcome non-target amplification of these two organisms by universal 18S primers, another set of blocking oligos were designed. We tested the oligos by preparing 18S amplicons from mock community templates (**Table S4**) containing bacterial, *A. thaliana*, *A. laibachii* and target *S. cerevisiae* genomic DNA. Quantification (qPCR) of target levels in the 18S libraries showed that blocking non-targets increased target levels between ~57x (1% target template) and ~57,000x (0.001% target template) (tested in the 18S V4-V5 region, **Fig. 4b**).

Here, we present the most complete picture of the *A. thaliana*-associated microbiome (and to our knowledge any microbiome) ever assembled in a single amplicon sequencing run (**Fig. 6**) based on combined diversity in 24 leaf samples collected from three wild locations. As expected, the 18S rRNA gene primers recovered a wide diversity of fungal and non-fungal eukaryotic microbiota, including various algae, cercozoa and amoebozoa (**Fig. 6a**). They even suggested that insects and helminthes are or were present on the leaves (**File S2**). The fungal and oomycete ITS datasets complemented the broader 18S data with more specificity in those groups – together, these two accounted for 44% of tree tips (observed genera, **Fig. 6a**). The prokaryote trees further demonstrate complementarity for primer sets targeting the same groups of microbes (**Fig. 6b**). Here, 42% of observed genera were discovered by both primer sets, with complementary diversity discovery especially in the phyla *Cyanobacteria* (V3-V4 dataset) and *Firmicutes* (V5-V7 dataset).

**Figure 6.**
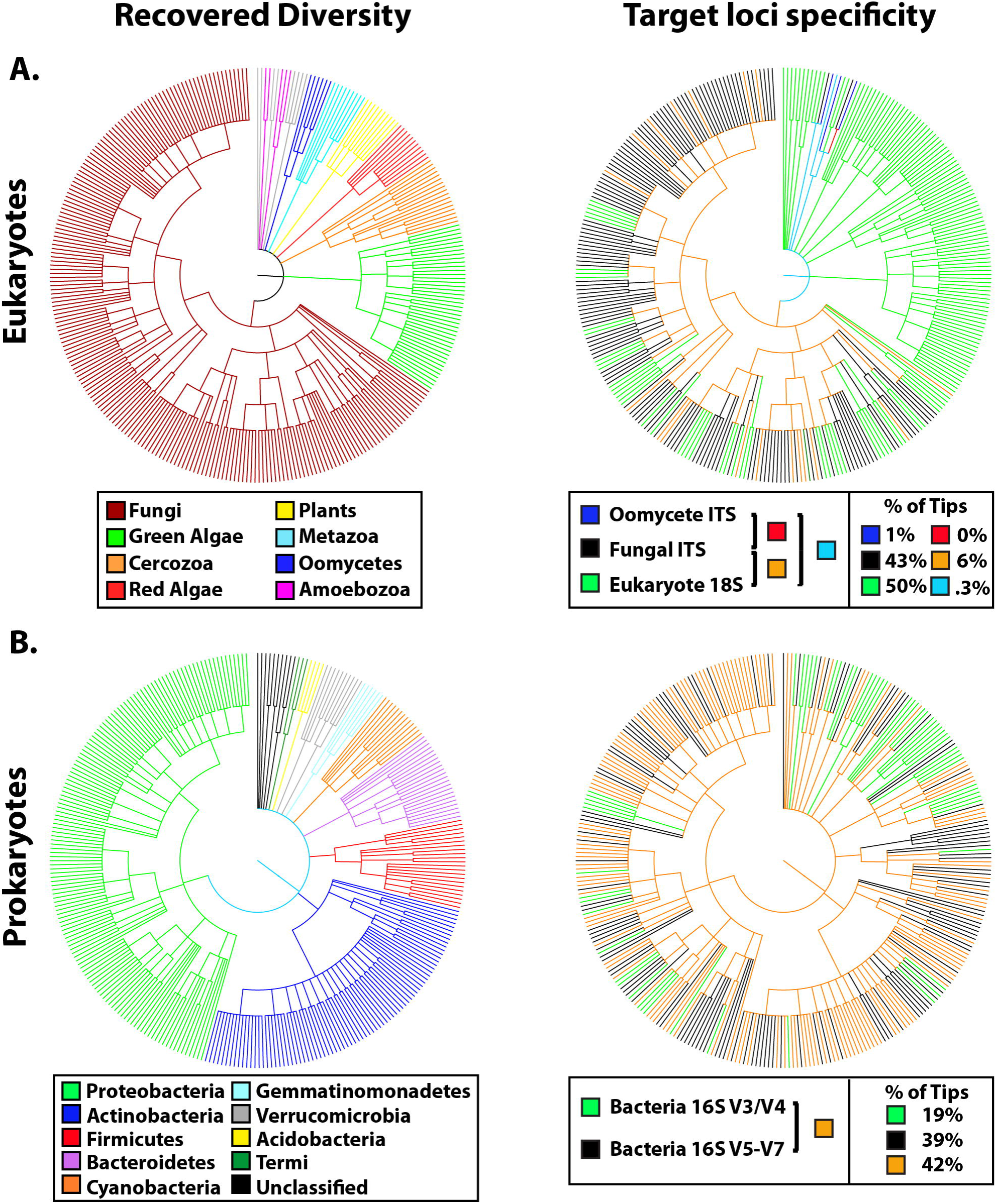
A comprehensive overview of high diversity microbiomes inhabiting *A. thaliana* leaves revealed by parallel amplicon sequencing of 8 loci targeting eukaryotes and prokaryotes microbes. A. 6 loci targeting eukaryotes: Two regions of the 18S rRNA gene (V4-V5 and V8-V9), two regions of the fungal ITS (ITS 1 and 2) and two regions of the oomycete ITS (ITS 1 and 2) revealed a diverse eukaryotic microbiota. The 18S loci revealed the broadest diversity but was complemented by fungi and oomycete-specific primer sets which had more detailed resolution within these groups. “Target loci specificity” refers to the taxa identified with each target group (Eukaryotes) or locus (Prokaryotes). B. 2 loci targeting prokaryotes: Two regions of the 16S rRNA gene (V3-V4 and V5-V7) that amplify mostly bacteria revealed a largely overlapping diversity profile complemented by unique discovery of taxa from each of the two target regions.

## Discussion

Amplicon sequencing of phylogenetically or functionally informative loci has become an indispensable technique in a variety of biology-related fields because its targeted approach (compared to untargeted approaches like metagenomics) allows the most accurate annotation possible by using specialized databases (DeSantis *et al.*, 2006). It has revealed that microbial community structuring is more complex than previously thought and suggested extensive interactions between (a)biotic factors and microbes (de Menezes *et al*, 2015) and between microbes even across kingdoms (Agler *et al*, 2016, Lima-Mendez *et al*, 2015). To understand these interactions, microbiome researchers need to be able to more completely characterize diversity in a single sequencing run. The current method enables this by targeting at least 8 loci in parallel. This drastic increase in resolution critically overcomes an inherent uncertainty in systems-scale investigations of factors contributing to microbial community structures.

A key technique enabling the current advances is employment of a two-step amplicon library preparation as opposed to a single step amplification. Many commonly used protocols (e.g., the Earth Microbiome Project 16S protocol based on (Caporaso *et al.*, 2011)) recommend using large, concatenated primers and one-step amplification. The advantages in technical reproducibility of a two-step approach that excludes concatenated primers in the first step were already described by Berry et al. (Berry *et al.*, 2011) for T-RFLP or 454 amplicon sequencing. For Illumina sequencing, concatenated primer bias was addressed with a 3-step approach: A 2-step amplification plus adapter ligation (Herbold *et al.*, 2015). That approach also allowed characterization of multiple gene regions, but a “head” sequence was concatenated to universal primers in the first amplification step. Our use of only universal primers in the first step, therefore, probably explains why mock community structure recovery was very accurate and replicable. Additionally, our approach adds all required adapters for sequencing in the second amplification step, eliminating problems associated with adapter ligation (Sambrook *et al.*, 1989).

Another major problem in amplicon sequencing is associated with using “universal” primers that in host-associated amplified non-target species, sacrificing read depth and masking diversity (Hanshew *et al.*, 2013). Previously, peptide nucleic acid “clamps” were used that were highly specific to non-target templates and which physically block their amplification (Lundberg *et al*, 2013). These clamps work efficiently in single-step amplifications, but their production is expensive, limiting rapid development and deployment of multiple clamps for new loci or for blocking several non-targets. Other approaches, like using oligonucleotide clamps that physically block the universal primer binding site (Vestheim & Jarman, 2008) are not applicable here because target and non-target binding sites are too highly conserved. Alternatively, blocking oligos are versatile and cheap and can be extensively tested at very low costs and adapted to virtually any target. Some universal primers targeting fungal ITS amplify fungal targets more efficiently than host, which explains why we had only minor host contamination in mixed mock community libraries. However, when fungal templates are less abundant they can significantly amplify plant ITS (Ihrmark *et al*, 2012). Because blocking oligos did not bias results, it is beneficial to always include them when relative abundance of target and non-target DNA is unknown.

One of the most persistent problems identified so far in amplicon sequencing is vast inflation of OTU diversity, mostly caused by sequencing errors (Kunin *et al*, 2010). Despite useful approaches to remove erroneous reads (Bokulich *et al.*, 2013, Reeder & Knight, 2010) or to reduce erroneous OTUs (Edgar, 2013) false diversity trends are commonly observed (Sinclair *et al.*, 2015). Errors explain the popularity of phylogenetics-based tools for beta diversity estimation based on the Kantorovich-Rubinstein metric, such as UniFrac (Evans & Matsen, 2012, Lozupone & Knight, 2005) or alpha diversity metrics like Faith’s PD (Faith, 1992). These weight differences in samples caused by distantly related OTUs more heavily since erroneous OTUs should be phylogenetically closely related. They have been very successful in identifying real differences between samples even when sequencing error is high. However, they are generally not applicable to loci like ITS, where extreme variability makes drawing phylogenetic relationships between all sequences questionable (Schoch *et al.*, 2012). Additionally, the assumptions of phylogenetic approaches do not hold when distantly related microbes occupy similar niches. For example, the basidiomycete yeast-like *Pseudozyma* spp. are phenotypically and ecologically much more similar to species like *Dioszegia* sp. (Inácio *et al.*, 2005) than to plant pathogenic members of its close relative *Ustilago* sp (Lefebvre *et al.*, 2013). Therefore, complementary approaches are needed that are sensitive to shifts in abundance among closely related taxa but which accurately delineate true and erroneous taxa.

The OPU approach addresses this problem because they are in principle like phylogenetic diversity metrics – very closely related (likely erroneous) OTUs are grouped into a unit which can be used to generate standard beta or alpha diversity metrics. Therefore, the results are less abstract than UniFrac or Faith’s PD (which lacks a taxonomic unit) and should be more sensitive to changes in abundance of closely related taxa. The term OPU was discussed elsewhere (Pernthaler & Amann, 2005) in the context of using phylogenetic grouping of organisms to move away from a specific percent identity as a working taxonomic unit but not as a systematic way to group erroneous OTUs. This concept was implemented in approaches to dynamically group amplicon reads by phylogeny based on tree cutting. Here clusters of reads were identified by training on a subset of data with known taxonomies (White *et al.*, 2010) or by known differences in substitution errors between or within species (Zhang *et al.*, 2013). General applicability of these approaches is unclear because of the major computational resources needed to cluster raw reads and because inferring phylogeny among all reads is questionable at some highly divergent loci like ITS (Schoch *et al*, 2012). Our implementation on the other hand uses pre-clustering of reads into OTUs and taxonomic groups. In this way large datasets are not a barrier because parallelization can be maximized according to available resources. Further, OTUs split by a taxonomic rank, e.g., family, are closely related and phylogenetic relationships can be determined even at highly divergent ITS loci. Therefore, diversity metrics based on OPUs represent a much needed phylogenetic method for loci that are not conserved enough to build alignments for example for UniFrac distances.

The realization of the immense complexity of biological systems – and our inability to adequately describe them - has led to many important, unresolved issues. For example, there is ongoing debate about what it means to view macroorganisms as holobionts, since symbiotic microbiota affect host health and fitness (Brucker & Bordenstein, 2013, Sharma *et al.*, 2014). Unanswered questions also linger, like what causes host genotype-independent taxonomic conservation of plant root microbiomes over broad geographic distances (Hacquard *et al.*, 2015). The tools described here will significantly increase the ability of researchers to accurately resolve microbial communities, addressing one of the primary limitations to progress. Although challenges remain, we expect this approach to equip researchers to make better hypotheses and to address seemingly intractable questions. These advances will thereby assist in increasing discovery of the important roles of microbiota.

## Acknowledgements

We wish to thank Ariane Kemen and Jonas Ruhe for providing *P. capsici* and *Pseudozyma* sp. isolates. We also thank the MPIPZ genome center for implementing our custom sequencing protocol. MA, EK, ND and SH were supported financially by the Max Planck Gesellschaft. AM, SH and ND were supported by the “Cluster of Excellence on Plant Sciences” program funded by the Deutsche Forschungsgemeinschaft. ND as also supported by a European Research Council advanced grant (ROOT MICROBIOTA) to Paul Schulze-Lefert.

## Author contribution

MA and EK conceptualized the project, designed and developed the methodology and wrote the manuscript. For the technology comparison, ND and SH performed experiments and MA, ND and SH analyzed the data. AM adapted the method to the 18S rRNA region and assessed plant-associated microbial communities. MA performed all other experiments and designed and wrote the scripts to process and analyze the data.

